# How Top-Down and Bottom-Up Regulation in Fronto-Amygdalar Network Changes over Time during Drug Cue-Exposure: An fMRI Study among Abstinent Heroin User

**DOI:** 10.1101/678961

**Authors:** Arash Zare-Sadeghi, Mohammad Ali Oghabian, Mehran Zare-Bidoky, Seyed Amir Hossein Batouli, Hamed Ekhtiari

## Abstract

Top-down regulation is one of the major neural cores in drug-craving management and relapse prevention. The dynamic temporal behavior of top-down regulation between the dorso-lateral and ventro-medial prefrontal cortices (DLPFC and VMPFC) and amygdala during drug cue-exposure has not been studied yet. Fifteen abstinent participants with heroin use disorder were scanned using drug cue-induced craving fMRI task. Using Dynamic Causal Modelling (DCM), the winning model showed a significant reciprocal connection between the VMPFC and DLPFC while there was a one-way effect of the VMPFC on the amygdala. There is also a top-down modulation by DLPFC on the VMPFC-Amygdala connection. Craving contrast input only modulated amygdala directly. Using sliding-window for temporal evaluation, craving input to amygdala increased over time, simultaneously, DLPFC top-down modulatory effect on VMPFC-amygdala connection decreased. Temporal changes in the network connectivity during cue exposure with enhancement in craving input to amygdala and reduction in top-down modulatory effects of DLPFC, could provide us with new insights towards the dynamic nature of the cue-reactivity and failure to control its motivational consequences. Dynamic response of top-down regulatory networks during cue exposure can be considered as a new potential biomarker in the future addiction fMRI studies.

## 1. Introduction

Cue-reactivity is a core phenomenon in the field of drug addiction and the extent of the patient’s response to salient cues can be an index for addiction severity and treatment success [1]. Pathologic response to salient cues -which might be either appetitive (seeking positive feelings), aversive (avoiding negative feelings) or both [2]-is a crucial neurocognitive process in people with substance use disorders (SUDs) and it is in accord with craving management, emotion regulation [3, 4] and self-control [2]. In addition to SUD, pathologic reactivity to some specific salient cues is the main symptom of many other addictive-type behaviors, like gaming disorders, pathological gambling or over-eating [5] and several mental health disorders, like post-traumatic stress disorder, social anxiety disorder and specific phobias [3]. Specifically talking about drug addiction, studies have shown that increased reactivity to drug-related salient cues is an important factor in inducing subjects (both humans and animals) to be susceptible to consume the cued substance and even relapse to drug use [2, 6].

The regions associated with emotion/motivation generation/regulation like the amygdala, striatum, thalamus, insula, and prefrontal cortex (PFC), are shown to have increased activity during cue-exposure in people with SUDs according to functional imaging studies [6-9]. Among the aforementioned regions, amygdala seems to be one of the determinant regions in cue-reactivity both in humans and animals [3, 4, 10]. As many studies have demonstrated, some of the main functions of the amygdala, regarding to salient cue-reactivity, are to generate, express, experience negative emotions [10], and aid in the process of conditioning for learning the rewarding value of cues [11].

Yet, amygdala doesn’t seem to be functioning independently during cue-reactivity and there are some cortical regions hypothesized to regulate amygdalar activities in various brain networks [12, 13]. One of the prominent networks for cue-reactivity processing is the Fronto-Amygdalar Network (FAN) [4, 12]. The PFC is involved in emotion regulation and executive functions, especially inhibitory control processes [14]. Some neurocognitive models of addiction suggest that hypersensitivity to drug cues in brain reward regions –including the amygdala-causes these regions to uncouple more easily from regions involved in top-down regulation located in the PFC and therefore, leads to seeking for drugs [2]. Many studies both on human and non-human subjects showed that the ventro-medial PFC (VMPFC) and amygdala are functionally and anatomically connected [4, 15]. The region presumed to influence the interactions between the VMPFC and amygdala is the dorso-lateral PFC (DLPFC) which its activity and its effective connectivity with the VMPFC are associated with better self-control and choosing delayed rewards [16, 17]. Both fMRI and brain stimulation studies have also pointed to the activation and causal role of prefrontal areas, especially the DLPFC during attempts to self-control and reducing drug craving [18]. Thus, the network including the DLPFC, VMPFC, and amygdala is supposed to be an important part of cue processing with a pivotal role in a person’s decision to exert cognitive control and successfully resist the temptation or to fail and seek drug use.

Although many studies showed the role of coupling between the PFC and amygdala in self-control, neuroimaging studies evaluating the temporal dimension of FAN connectivity are scarce in the field of addiction. According to the psychological background of the effect of temporality, Baumeister and his colleagues declared a “strength” model for self-regulation [19]. Despite the controversies on this model among psychologists [20], there are many studies consistent with the strength model, demonstrating that the more the subjects are doing inhibitory or memory tasks, the less likely they are in control for their food or drug craving afterwards [20]. From a neuroimaging perspective as well, persistent with time-variable changes, Wagner and his colleagues in their fMRI study, where participants tried to self-regulate after a hard cognitive task, showed attenuated amygdalar activity, implying a probable underlying mechanism for resource depletion in accordance with the strength model [4].

The aim of our fMRI study was to assess the effective connectivity between the amygdala, VMPFC, and DLPFC within the FAN during drug cue-exposure and its temporal dynamism. We implemented Dynamic Causal Modeling (DCM) analysis combined with fMRI data to study the effective connectivity in FAN. By using sliding temporal windows, we also evaluated the effect of time on FAN connectivity. To the best of our knowledge, this is the first study focused on the effective connectivity and temporal changes in FAN during cue-exposure among people with SUDs.

## 2. Methods

### 2.1. Participants

This study was approved by the “*independent ethical committee”* of Tehran University of Medical Sciences. Eighteen right-handed (the right-handedness was assessed using the Edinburgh Inventory [21]) male subjects with heroin use disorder were recruited for this study. They all have been successfully abstinent after an extended detoxification program. Subjects were all crystalline-heroin smokers before their treatment initiation in a program affiliated to a non-governmental not-for-profit organization named “*Congress 60*”, Tehran, Iran. They also met heroin use disorder criteria for at least 6 months before their treatment, based on the Diagnostic and Statistical Manual of Mental Disorders (DSM V). All subjects had negative results in urine tests for opiates, stimulants, and benzodiazepines for at least 30 days before the day of scanning. In the scanning day, Tobacco smoking was not restricted. We excluded subjects with any history of head trauma, major medical or psychiatric disorders (other than heroin use disorder with or without other SUDs), and visual field deficiency. We also excluded 3 subjects’ data; 1 for structural image problem (the MPRAGE image was damaged), and 2 for excessive motion in EPI images. The age range of the remaining 15 subjects was 20-35 years with a mean of 29. The detailed participant specifications can be found in Table 1. All subjects were provided with written informed consents to participate in this study.

**Table 1:** The demographic and substance use profile of participants (n=15).

|  | Descriptive Statistics |
| --- | --- |
| Gender (male) | 15/15 |
| Age | 28.18 ± 1.93 |
| Education | 11.21 ± 1.16 |
| Marital Status (married) | 10/15 |
| Duration of opiate abstinence (month) | 13.56 ± 3.57 |
| Subjects with heroin use disorder | 15/15 |
| Duration of crystalline heroin use disorder (years) | 1.92 ± 1.2 |
| Subjects with opium use disorder | 15/15 |
| Duration of opium use disorder (years) | 6.97 ± 2.1 |
| Subjects with brown heroin use disorder | 5/15 |
| Subjects with alcohol use disorder | 12/15 |
| Subjects with hashish use disorder | 14/15 |
| Subjects with stimulant use disorder | 4/15 |
| Subjects with cigarette smoking | 15/15 |

### 2.2. FMRI task

The task was a visual block design with 24 heroin-related images and 24 neutral images; the drug images were selected from a previously validated database for people with heroin use disorder [22] and 24 neutral stimuli that were selected from International Affective Picture System [23] and matched on the basis of the object category class and visual features. There were 6 runs in the task; each run consisted of four blocks in a fixed order: 24s rest (a fixation cross was shown), 24s neutral (4 neutral images were shown, each for 6s), 24s rest, and 24s craving (4 heroin-related images were shown, each for 6s). The task length was about 10 minutes. We have introduced this fMRI task in our previous publications [9]. The task was projected on a white screen in front of the scanner and viewed by a mirror system mounted on the head coil. The details of the task are depicted in Figure 1.

**Figure 1:**
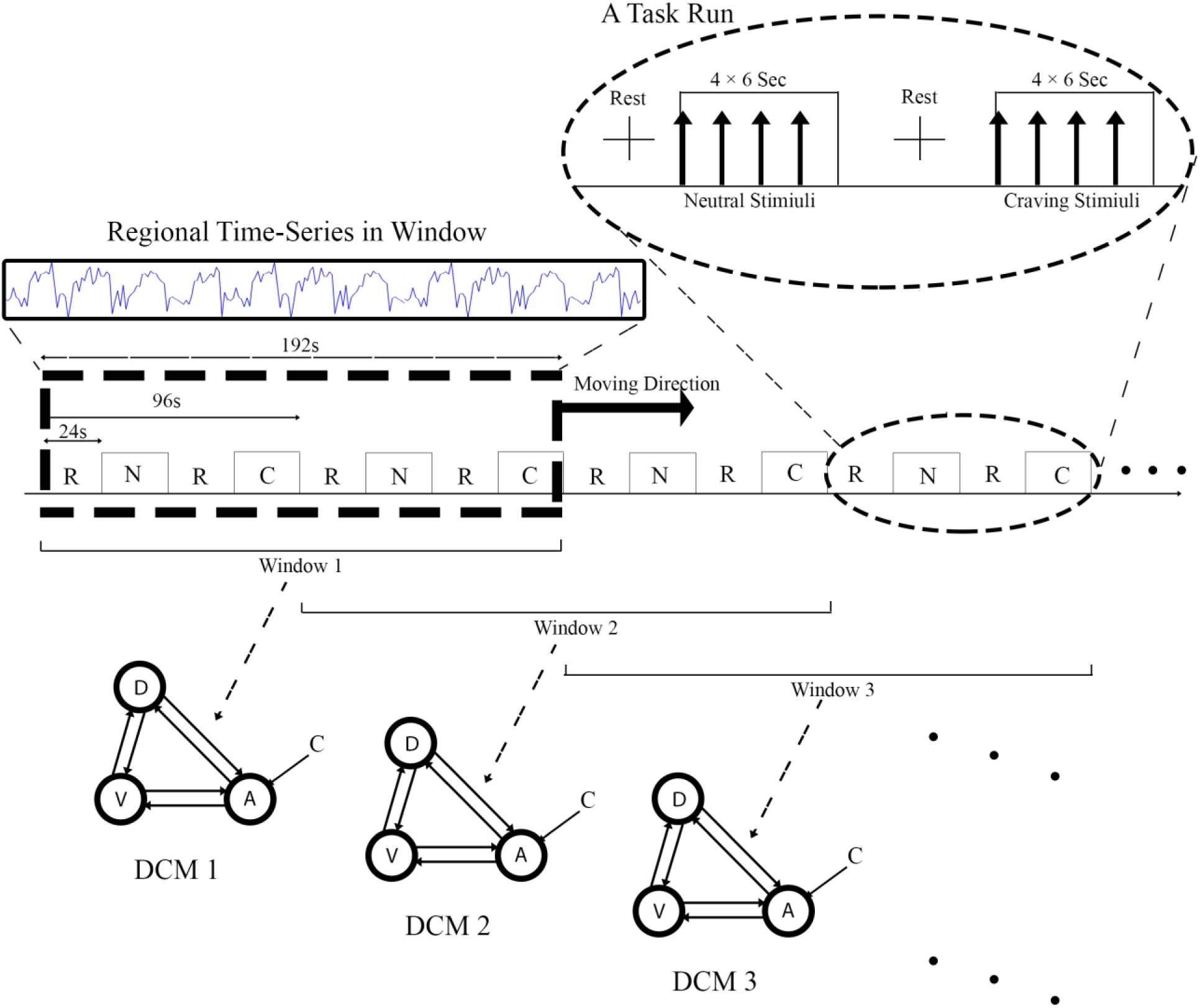
The schematic of the fMRI task and the Sliding Window DCM. As it is shown in the top-right of the figure, each run of the task included two rest blocks in which a cross was shown to the subject, a Neutral block in which four neutral images were shown to the subject each for 6 seconds, and a craving block in which four heroin images were shown to the subjects with the same duration as neutral images. Each block was 24 seconds, so the length of each run was 96 seconds. The task had six runs and the total duration was 576 seconds. As it is shown in the top left of the figure the window size was as long as two runs or 192 seconds and it was moved 96 seconds in each step. The time-series of each five resulted parts was extracted and was used for estimating the related effective connectivity model (DCM).

### 2.3. Data acquisition

Functional and Structural images were acquired using Avento 1.5T scanner; Siemens, Germany. T1-weighted images had the following properties: MPRAGE, TE= 3.55ms, TR= 1910, voxel size = 1 × 1 × 1mm^3^, and flip angle = 30 degrees. T2*-weighted images had the following properties: TR= 3000, TE= 50ms, matrix size = 64 × 64, voxel size = 3 × 3 × 3 mm^3^, and flip angle 90 degrees.

### 2.4. Data analysis

#### 2.4.1. Preprocessing

The software package FSL5 (FMRIB’s Software Library, www.fmrib.ox.ac.uk/fsl) [24] was used as a preprocessing tool for removing any unwanted parts of the data. The motion correction was done using the MCFLIRT [25] tool with 6 degrees of freedom, next the slice timing correction was done with interleaved order, then a Gaussian kernel with full width at half maximum (FWHM) equal to 5mm was used to smooth the data. Using the FLIRT tool [25], the echo planar images first were registered to the structural image with 12 degrees of freedom and then the affine transformation was used to register them to the MNI space. High-pass filtering was done using a filter with the cut-off frequency equal to the inverse of 96 seconds; this is the length of one block. All EPI images were intensity normalized, so the group analysis could be done in the next steps.

#### 2.4.2. Activation analysis

Three ROIs were selected according to the a priori hypothesis of the study; the VMPFC, DLPFC, and amygdala. We used masks for each region, which were all in MNI space and to register the mask to each subject’s images, the transformation matrices, derived from the registration step of preprocessing, were used. Using this method, each subjects’ data had their own specific masks.

The FEAT tool [26] from the FMRIB’s package was used to model the data and find the activations in the brain. Canonical hemodynamic response with its derivative was used to model the regressors for two conditions of interest; craving and neutral. Next, an ROI based analysis was carried out using the FLAME2 [27] tool of the FSL5 package, as the group level analysis for the contrast “craving > neutral”.

#### 2.4.3. Time-series extraction

The eigen-variate tool of SPM12 (Statistical Parametric Mapping; Wellcome Department of Cognitive Neurology, London, UK, https://www.fil.ion.ucl.ac.uk/spm/) was used for extracting each region’s time-series. Three masks, one for each region, were defined in MNI space. For each subject, the transformation matrix between the MNI space and EPI space was used to map the mask to each subject’s own functional space. This method gave us the best registration result but it was depending on the masks’ initial definition. To get the best results we checked the accuracy of the initial masks with an expert anatomist. We did not threshold the voxels included in each region for any contrasts; so all voxels in the regions had their effects on the final extracted time-series. Using limited voxels may result in stronger connections in the final effective connectivity network, but we preferred to use the raw time-series and omitted the possibility of imposing any unwanted bias on the results.

#### 2.4.4. Effective connectivity

DCM was used for quantifying effective connectivity network. DCM needs some steps to reach an acceptable network. The first step of DCM is defining a model space which has the following properties: first, it cannot have all possible models because the process of estimation will take too long; second, its space must include all possible models which are meaningful from a neuroscience perspective; and third, its model space may include some other new models for testing new hypothesis [28]. We included 16 models in our model space. In our a priori hypothesis we were interested in the modulatory effect of the DLPFC on the link from the VMPFC to the amygdala, so we included models with and without this effect. We used DCM12 tool integrated in SPM12 to specify the model space and then estimate all models for each subject.

Bayesian Model Selection (BMS) is a part of DCM algorithm which compares estimated models and checks complication and fitness of the models both at the same time. The BMS algorithm output is the exceedance probability of the models. We first implemented BMS on single models, without considering our hypothesis and then -using family partitioning- we tested our hypothesis for the DLPFC effect. Models with this effect built one of the families and other models built the other ones. Bayesian Model Averaging was used on the winning family of the previous step to reach the final effective connectivity network.

#### 2.4.5. Time variability analysis

The nature of brain effective connectivity networks are time variable and this has been proved in many neuroimaging modalities [29-31]. We have used DCM for fMRI data to reach the effective connectivity network. DCM is based on dynamic changes, but the final concluding networks of DCM algorithm do not quantify dynamic changes during the time. To show the time variability, we have used a sliding window on the data. The window size was as long as two runs (192 seconds) and the step size for moving the window was chosen to be one run (96 seconds) ended up to extract 5 time-series. Using these extracted time series, we had five independent DCMs to estimate connectivities for each subject’s whole time series. This is depicted in Figure 1. We considered each DCM as a time point; thus, the changes between those networks were assumed to be the changes during the time. In this step, we did not include the model space and only re-estimated the parameters of the winning network, which had been quantified in the classic DCM step.

## 3. Results

### 3.1. ROI based activation analysis

The contrast of interest in this study was the difference of the activation patterns when the subjects were shown the craving-inducing pictures versus the neutral pictures (Craving > Neutral). The activation map of this contrast is depicted in Figure 2 for ROI analysis while Table 2 contains the results for whole brain analysis. The ROI analysis results were corrected voxel-wise and the whole brain analysis results were corrected cluster-wise with Family Wise Error Rate (FWER) method (p-value <0.05).

**Table 2:** The whole brain analysis results. Six clusters were found to have more than 200 voxels. The results were corrected cluster-wise with FWER method.

| Anatomical Regions | Cluster Size | Z-values | Local Maximums Co-ordinates (MNI) |  |  |
| --- | --- | --- | --- | --- | --- |
| Supracalcarine Cortex | 3123 | 4.07 | 0 | -90 | 8 |
| Lingual Gyrus (R) |  | 3.96 | 2 | -84 | 6 |
| Lingual Gyrus (L) |  | 3.93 | -12 | -76 | -8 |
| Cuneus (L) |  | 3.91 | -10 | -98 | 12 |
| Retrosplenial cortex (R) |  | 3.66 | 12 | -50 | 4 |
| Cuneus (R) |  | 3.61 | 8 | -76 | 28 |
| Superior parietal lobule (L) | 1228 | 3.57 | -2 | -40 | 66 |
| Primary somatosensory cortex (R) |  | 3.46 | 8 | -44 | 68 |
| Superior parietal lobule (L) |  | 3.44 | -12 | -40 | 52 |
| Superior parietal lobule (L) |  | 3.41 | -18 | -42 | 42 |
| Primary somatosensory cortex |  | 3.29 | 0 | -26 | 62 |
| Superior Parietal Lobule (R) |  | 3.28 | 14 | -48 | 72 |
| Parahippocampal Gyrus (R) | 314 | 3.73 | 26 | -44 | -8 |
| Fusiform Cortex (R) |  | 3.42 | 36 | -46 | -10 |
| Parahippocampal Gyrus (R) |  | 2.84 | 38 | -26 | -14 |
| Fusiform Cortex (R) |  | 2.58 | 38 | -30 | -16 |
| Primary motor cortex (R) | 293 | 3.68 | 20 | -18 | 66 |
| Primary motor cortex (R) |  | 3.52 | 32 | -28 | 62 |
| Primary motor cortex (R) |  | 3.46 | 36 | -30 | 66 |
| Primary motor cortex (R) |  | 3.31 | 30 | -28 | 68 |
| Primary motor cortex (R) |  | 2.75 | 12 | -28 | 72 |
| Parahippocampal Gyrus (L) | 258 | 3.51 | -20 | -42 | -10 |
| Fusiform Cortex (L) |  | 3.43 | -28 | -48 | -6 |
| Heschl's Gyrus (R) | 226 | 3.48 | 54 | -26 | 18 |
| Secondary somatosensory cortex (R) |  | 3.36 | 52 | -28 | 24 |
| Heschl's Gyrus (R) |  | 2.85 | 62 | -24 | 12 |
| Secondary somatosensory cortex (R) |  | 2.73 | 40 | -28 | 28 |
| Supramarginal Gyrus (R) |  | 2.61 | 48 | -38 | 14 |
| Angular Gyrus (R) |  | 2.46 | 58 | -36 | 18 |

**Figure 2:**
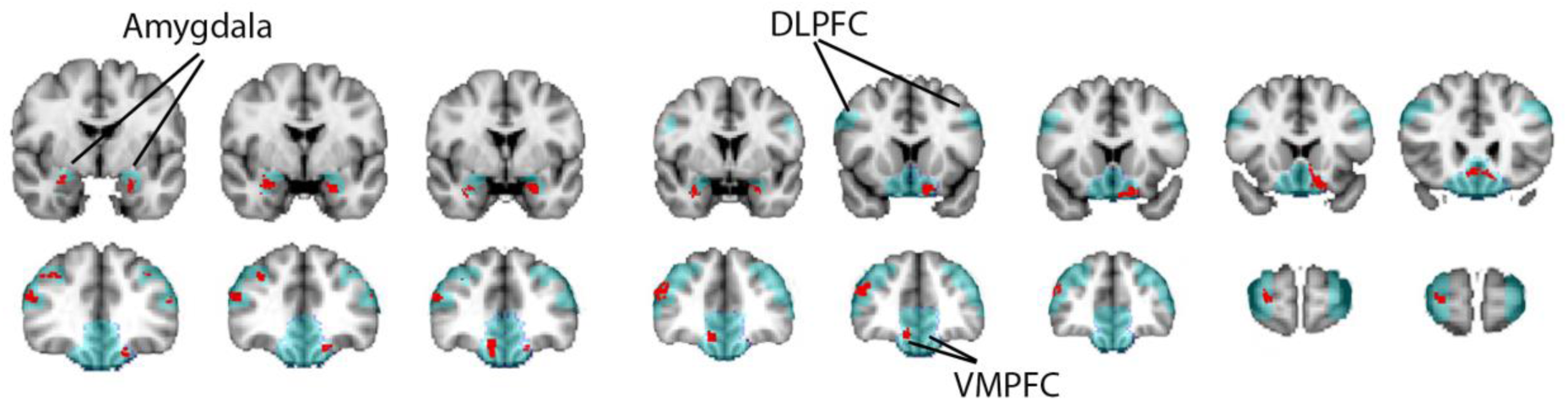
The result of ROI analysis (coronal view). Three regions of interest were selected; Amygdala, DLPFC, and VMPFC. Blue colors show these three regions and red regions are activated parts of these three.

### 3.2. Time-invariant DCM

We first implemented Bayesian Model Selection (BMS) on the estimated models without any family partitioning. One model was removed from the model space due to its low BMS probability result. Next, as mentioned before, we divided the model space into two families (with/without the DLPFC effect on the VMPFC and amygdala link) and implemented BMS on these families. The results showed the dominance of models with the effect of the DLPFC on the link from the VMPFC to the amygdala as depicted in Figure 3.

**Figure 3:**
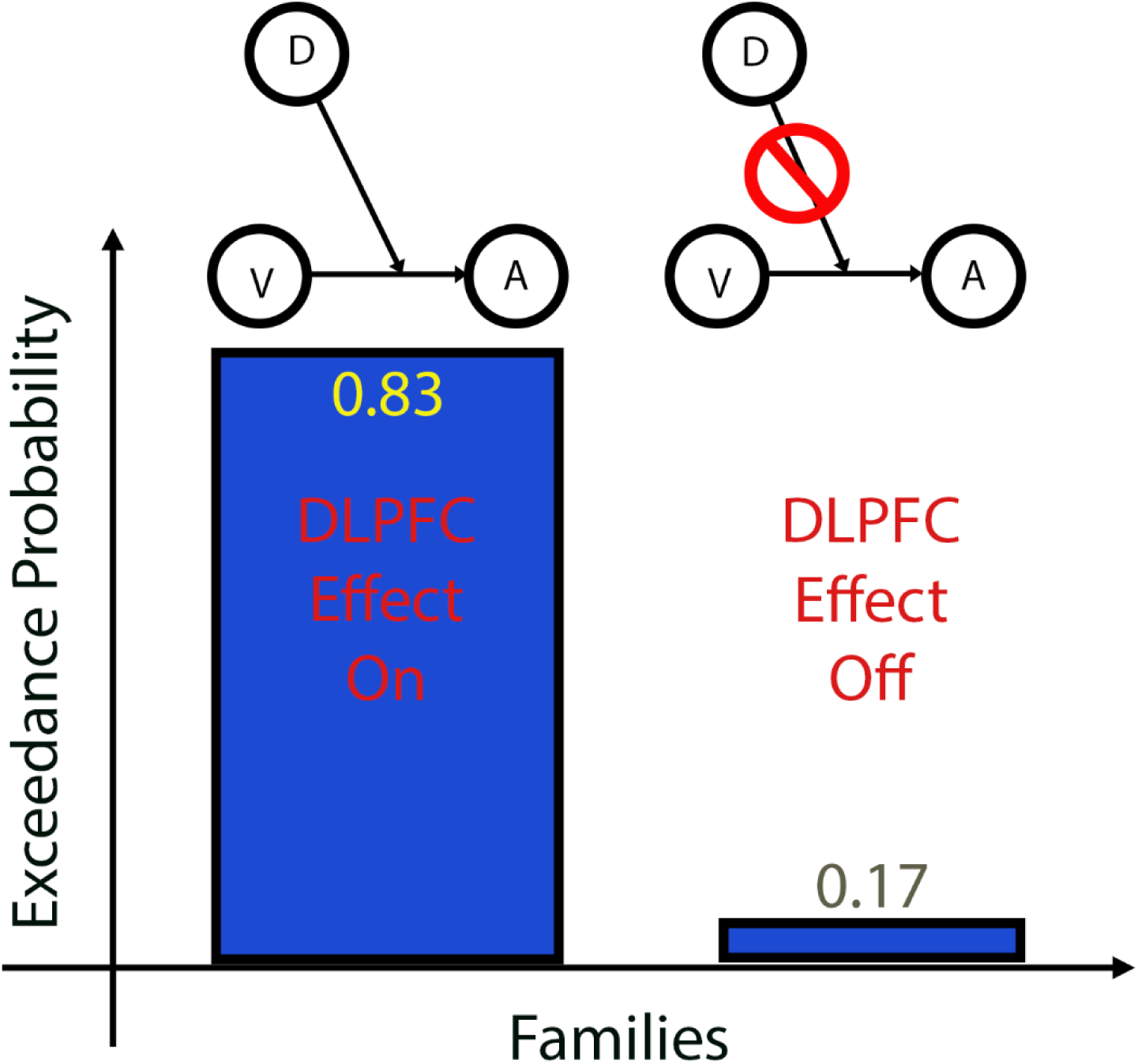
The Bayesian Model Selection (BMS) results for the assigned two families. The first family (left) includes models in which DLPFC has an effect on the link from the VMPFC to the amygdala and the second family (right) includes models in which the effect does not exist. The vertical axis shows the exceedance probability value of these families.

Bayesian Model Averaging (BMA) was implemented on the winning family of the previous step and the result was a DCM network as is depicted in Figure 4 (top panel). All three regions were connected to each other but the connection between the VMPFC and amygdala was not reciprocal and was only from the VMPFC to the amygdala. Self-inhibitory effects were seen in all regions and the craving contrast input had a significant effect only on the amygdala. Our hypothesis about the DLPFC effect was proved and the link’s strength was significant.

**Figure 4:**
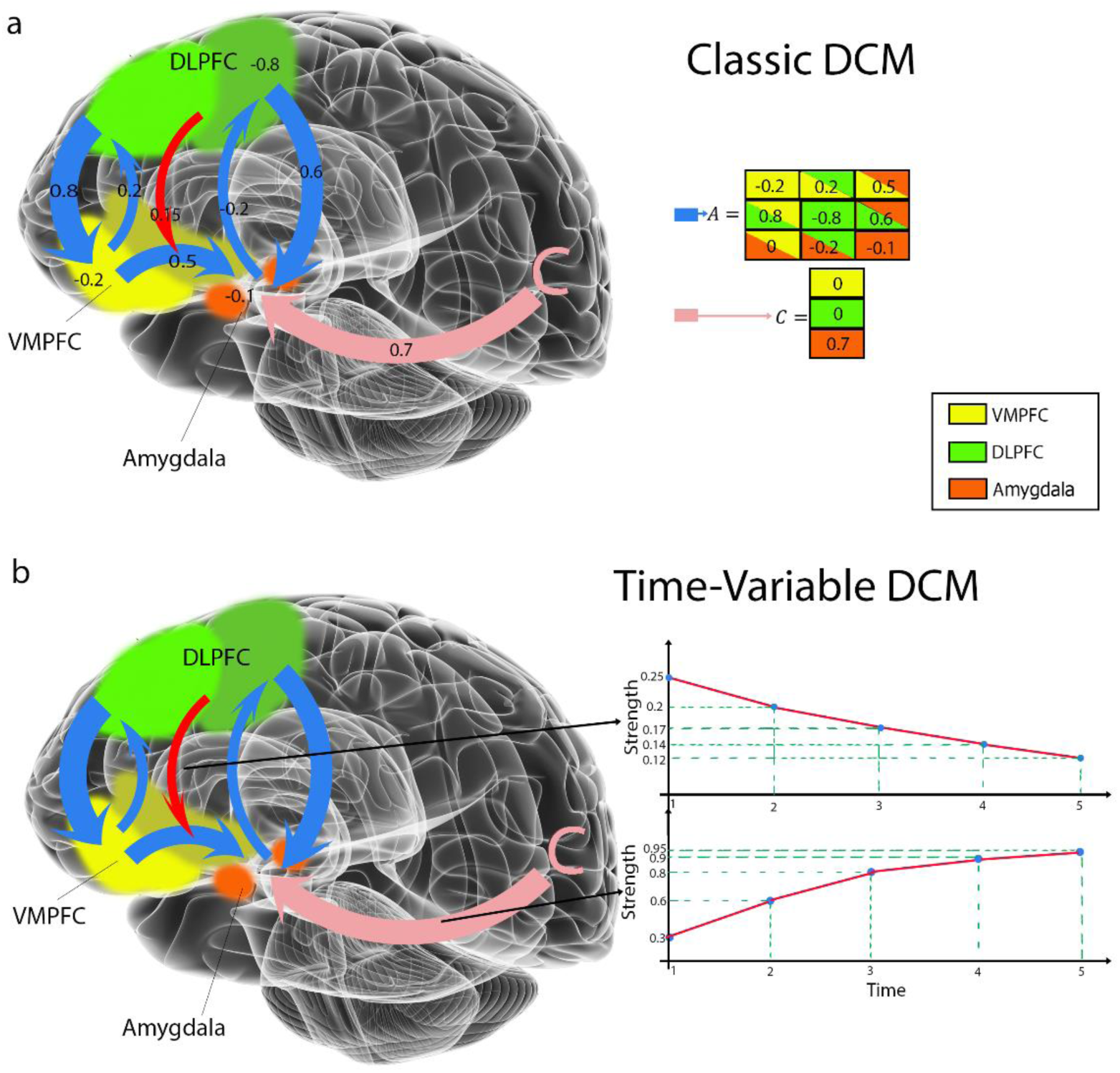
The classic and time-variable DCMs in Fronto-Amygdalar Network. Three Regions of Interest (ROIs) are shown; DLPFC (green), VMPFC (yellow), and Amygdala (orange). The connections between ROIs are shown in blue arrows, the effect of the DLPFC on the link between VMPFC to the amygdala is shown in red arrow, and the craving contrast input link is shown in pink arrow. (a) The classic DCM results and links are shown in the upper panel. (b) The time-variable DCM results are shown in the lower panel; The two time-variable links’ strengths are shown in the diagrams. The horizontal axis is time and the vertical axis is the link’s strength. The decrease in the strength of the effect of the DLPFC on the link between VMPFC to the amygdala is shown in the top diagram (Strength= 0.25(1-exp(2.6t)) and the increase in the strength of craving contrast input link is shown in the bottom diagram (Strength= 0.65exp(2.16t).

### 3.3. Time-variant DCM

The concluded network of the previous step was used in the next step. Using a 192s sliding window, moving for 96s in each step, we extracted five fMRI time series. Five different DCM networks with the same structure were re-estimated, accordingly. Parameters of this network or the links’ strength changed each time the estimation process was implemented. The strength of craving contrast input to the amygdala increased as the strength of the DLPFC effect on the link from the VMPFC to the amygdala decreased. An exponential graph was fitted to each result and the time-constants of such exponential increase/decrease was calculated. The Figure 4 (bottom panel) depicts the changes of these two links’ strength.

## 4. Discussion

In this study, we examined the influence of drug cue-exposure over time on top-down and bottom-up regulation in the fronto-amygdalar network (FAN) among people with heroin use disorder for both static and dynamic dimensions of effective connectivity. The static DCM results posited a reciprocal connection between the DLPFC and VMPFC and a one-direction connection, from the VMPFC to the amygdala with the effect of DLPFC on this connection. As hypothesized, there were changes in the interactions between the DLPFC, VMPFC, and amygdala during cue-exposure. According to our DCM results, it is demonstrated that when subjects are being exposed to drug cues, craving input to the amygdala increases over time. Furthermore, the more the subjects are exposed to the drug cues, the more attenuation is seen on the top-down modulatory effect of the DLPFC on the VMPFC-amygdala connection.

### 4.1. ROI activations

By analyzing the fMRI data, participants showed increased amygdalar activity when they were exposed to drug cues compared to neutral pictures. In line with other studies, it is deduced that there is a relation between the higher desire for drug use and higher amygdalar activation [6, 7]. The higher activation of DLPFC when watching drug cues compared to controls is also supported by the controlling role of DLPFC on regulating the emotions [32] given that drug craving is also a highly emotional process. Augmented activation of VMPFC when watching the salient cue can be due to being involved in the process of reward valuing [33].

### 4.2. Static (Classic) FAN

In line with the results of the current study regarding the effective connectivity between the VMPFC, DLPFC and amygdala, DCM analysis in PTSD patients has shown a dominant connectivity from the amygdala to VMPFC [34] and also a reciprocal connectivity between the amygdala and DLPFC [35]. Modulating effect of the DLPFC on the VMPFC-amygdala connection was also observed in our study. This result was consistent with the significant role of the DLPFC in regulating the amygdalar activity during craving through its interaction with the VMPFC [16, 17]

### 4.3. Time-Variant FAN

According to our time-variant results, the more the subjects were exposed to the drug cues, the more decrease in the effect of DLPFC on the link between the VMPFC and amygdala was detected. Meanwhile, the strength of craving input to the amygdala increased over time. Given the decrease of the top-down regulation, and on the other hand increase of the bottom-top regulation, we can speculate that both contribute to bringing about an exaggerated response to salient drug cues with lack of control and increased drug craving over time during cue exposure.

One group of networks which overlaps with FAN in both anatomy and functions is the triple network model between Default Mode Network (DMN), Salient Network (SN), and Executive Control Network (ECN). The most prominent components of DMN and ECN are the medial PFC and lateral PFC respectively. DMN is responsible for interoception and self-oriented processes such as thinking about drugs [36]. ECN serves executive functions and causes goal-directed cognition [36] and its hypo-activity and dominance of DMN on it, can lead to relapse to drug use [37]. As some addiction theories suggest, addiction could be one of the sequelae of the deficit in executive function and cognitive control of SUD patients [38]. SN identifies which network fits best to the current condition and shifts dominancy to that network; the main regions suggested to take part in SN is the insula and amygdala [39-41]. It is reported that increased connectivity between SN to ECN can reduce withdrawal symptoms [42]. The imbalance between these networks also results in many behavioral and cognitive problems in both healthy subjects and people with SUDs [43]. As Li et al. demonstrated more vulnerability to relapse for subjects who their DMN is more active than ECN [44]. In accord with the results of our current study, we have previously shown in our study that higher levels of craving for drug use are associated with dynamic transition of functional connectivity between DMN-ECN to higher functional connectivity between SN-DMN [45]. The results of our current study also somehow can show the same results as the effects of the DLPFC (part of the ECN) on the connection between the VMPFC and amygdala attenuates which results in the dominance of SN-DMN.

By reviewing major psychological theories considering the limitation of self-control [20], these models have a core in common and that is “time-variability” of self-control. According to the “strength” model advanced by Baumeister and his colleagues, temporal processing limit would be demonstrated when engaging in a self-control task depletes the person’s resources of self-control [19]. Another major theory, called “process” model, advanced by Inzlicht et al., explains self-control as the salience of motivation to extrinsic rewards (“have to” task) and intrinsic rewards (“want to” task). When the salient motivation shifts from “have to” task to “want to” task, the person tends to quit the exploiting task and seek leisure [20]. The neuroscientific explanation provided by these models regarding the results of the present study can be that the increase in craving inputs to the amygdala (bottom-top augmentation) and decrease in the effect of the DLPFC on the VMPFC - amygdalar connection (top-down attenuation) that might be inferred either depletion, in the opinion of *the strength* model, or a shift in motivation, in the opinion of the *process* model.

### 4.4. Potentials for dynamic measures of cue reactivity to be validated as biomarkers

Neural activation patterns in patients with SUDs could provide us with insight into the severity of the disorder and other behavioral problems associated with it [46]. These findings could be biomarkers for the progression of the disease, treatment effectiveness and relapse prediction as validated by fMRI studies [47]. For instance, Janes et al. assessed activation of response to smoking-related cues in abstinent Tobacco smokers and demonstrated that the higher activation in the insula and PFC (including DLPFC) was associated with greater probability of relapse [48]. Also, another study showed that smokers who have more active lateral PFC during simple inhibition tasks show less cigarette craving in real life [49]. To conclude, we assume that the more the effect of DLPFC on the connection between the VMPFC-amygdala during cue-exposure remains, the more successful the subject in controlling the craving would be. The supporting notion for this idea is the study done by Janes et al. showing that people who failed to quit smoking had less functional connectivity between the amygdala and DLPFC at their pre-quit state [48]. As It is suggested, although the DLPFC activation could stay the same or even increase, the underlying reason for relapse could be the decrease in the effect of the DLPFC on the amygdala. There are still very few published brain imaging studies using predictive modeling to validate biomarkers for addiction medicine, however, all of them are using static markers. Future studies can see if there is a prediction power for the dynamic markers like the exponential slope of the attenuation in the effect of DLPFC on the VMPFC-amygdala connection in the individual level to explain variability in treatment outcome.

### 4.5. Limitations

Our fMRI task could potentially provide a better platform for exploring the temporal behavior of different networks during cue reactivity if it was longer and had more runs. The positive benefit of adding more runs to the task can be explored in the future. Furthermore, the small sample size limited our study to an initial exploration. Further studies with more subjects are needed to replicate these results. Our subjects have been abstinent for more than one year on average. Many studies have shown that different stages of abstinence can affect brain activations in response to drug cues [9, 50]. Also, among the other important factors are the effect of age and sex which explain some inconsistencies among the results in different studies [51], therefore, we should be careful in extending our results to other populations [52]. Exploring these results in subjects in other stages of substance use and recovery can provide a better picture of this phenomenon.

### 4.6. Conclusion

Most of the studies in the field of cue-reactivity are concerned with static brain activities, but this study for the first time addressed the time-variability of cue-reactivity in the FAN. Using network analysis by time-variant DCM, we report a gradual decrease in the DLPFC top-down regulation over VMPFC-amygdala connection and also a gradual increase in the craving input to amygdala as the exposure to salient drug cues lasts longer. These results help shed light on the dynamic nature of the cue reactivity and could give us a new intuition of how failure to control drug craving can happen in the brain. Future studies are required to investigate these results for different kinds of drugs, different brain networks and also different stages of recovery.

## Acknowledgments

Authors would like to thank Mr. Hossein Dezhakam, founder of the “Congress 60” NGO and its recovery community for their active contributions. We also like to thank Milad Kassaei and Arshiya Sangchooli for their invaluable comments and ideas through the process of the manuscript preparation. This study is supported by Tehran University of Medical Sciences by grants to MAO and HE.

## Authors Contribution

HE had the original idea of the manuscript and was supervising the process of the study from the start to end. AZS and AHB, has analyzed the fMRI data and implemented classic/time-variable DCM on the data to reach the quantitative network, he has also written the *Materials and Method* and *Results* parts of the manuscript with the help of SAHB. MAO, as the head of Neuro-Imaging and Analysis Group (NIAG), and HE had guided the research to reach its goal. HE and MZB drafted the manuscript and provided critical revisions of the manuscript for important intellectual content. All authors critically reviewed the content and approved final version for publication.

## Data Availability Statement

The datasets generated and analyzed during the current study are available from the corresponding author on reasonable request.

## References

1. Jasinska, A.J., et al., Factors modulating neural reactivity to drug cues in addiction: a survey of human neuroimaging studies. Neuroscience & Biobehavioral Reviews, 2014. 38: p. 1–16.

2. Heatherton, T.F. and D.D. Wagner, Cognitive neuroscience of self-regulation failure. Trends Cogn Sci, 2011. 15(3): p. 132–9.

3. Peters, J., P.W. Kalivas, and G.J. Quirk, Extinction circuits for fear and addiction overlap in prefrontal cortex. Learn Mem, 2009. 16(5): p. 279–88.

4. Wagner, D.D. and T.F. Heatherton, Self-regulatory depletion increases emotional reactivity in the amygdala. Soc Cogn Affect Neurosci, 2013. 8(4): p. 410–7.

5. Boutelle, K.N., et al., Increased brain response to appetitive tastes in the insula and amygdala in obese compared with healthy weight children when sated. Int J Obes (Lond), 2015. 39(4): p. 620–8.

6. Goudriaan, A.E., et al., Brain activation patterns associated with cue reactivity and craving in abstinent problem gamblers, heavy smokers and healthy controls: an fMRI study. Addiction biology, 2010. 15(4): p. 491–503.

7. Etkin, A. and T.D. Wager, Functional neuroimaging of anxiety: a meta-analysis of emotional processing in PTSD, social anxiety disorder, and specific phobia. Am J Psychiatry, 2007. 164(10): p. 1476–88.

8. Hassani-Abharian, P., et al., Exploring Neural Correlates of Different Dimensions in Drug Craving Self-Reports among Heroin Dependents. Basic Clin Neurosci, 2015. 6(4): p. 271–84.

9. Tabatabaei-Jafari, H., et al., Patterns of brain activation during craving in heroin dependents successfully treated by methadone maintenance and abstinence-based treatments. J Addict Med, 2014. 8(2): p. 123–9.

10. Banks, S.J., et al., Amygdala-frontal connectivity during emotion regulation. Soc Cogn Affect Neurosci, 2007. 2(4): p. 303–12.

11. Robbins, T.W., K.D. Ersche, and B.J. Everitt, Drug addiction and the memory systems of the brain. Ann N Y Acad Sci, 2008. 1141: p. 1–21.

12. Gold, A.L., R.A. Morey, and G. McCarthy, Amygdala–prefrontal cortex functional connectivity during threat-induced anxiety and goal distraction. Biological psychiatry, 2015. 77(4): p. 394–403.

13. Shou, H., et al., Cognitive behavioral therapy increases amygdala connectivity with the cognitive control network in both MDD and PTSD. Neuroimage Clin, 2017. 14: p. 464–470.

14. Everitt, B.J. and T.W. Robbins, Neural systems of reinforcement for drug addiction: from actions to habits to compulsion. Nat Neurosci, 2005. 8(11): p. 1481–9.

15. Zhang, S., et al., Ventromedial prefrontal cortex and the regulation of physiological arousal. Soc Cogn Affect Neurosci, 2014. 9(7): p. 900–8.

16. Hare, T.A., C.F. Camerer, and A. Rangel, Self-control in decision-making involves modulation of the vmPFC valuation system. Science, 2009. 324(5927): p. 646–8.

17. McClure, S.M., et al., Separate neural systems value immediate and delayed monetary rewards. Science, 2004. 306(5695): p. 503–7.

18. Shahbabaie, A., et al., State dependent effect of transcranial direct current stimulation (tDCS) on methamphetamine craving. International Journal of Neuropsychopharmacology, 2014. 17(10): p. 1591–1598.

19. Baumeister, R.F. and T.F. Heatherton, Self-regulation failure: An overview. Psychological inquiry, 1996. 7(1): p. 1–15.

20. Inzlicht, M., B.J. Schmeichel, and C.N. Macrae, Why self-control seems (but may not be) limited. Trends Cogn Sci, 2014. 18(3): p. 127–33.

21. Oldfield, R.C., The assessment and analysis of handedness: the Edinburgh inventory. Neuropsychologia, 1971. 9(1): p. 97–113.

22. Ekhtiari, H., et al., Designing and Evaluation of Reliability and Validity of Five Visual Cue-induced Craving Tasks for Different Groups of Opiate Abusers. Iranian Journal of Psychiatry and Clinical Psychology, 2008. 14(3): p. 337–349.

23. Lang, P.J., M.M. Bradley, and B.N. Cuthbert, International affective picture system (IAPS) : affective ratings of pictures and instruction manual. 2005, Gainesville, Fla.: NIMH, Center for the Study of Emotion & Attention.

24. Jenkinson, M., et al., FSL. Neuroimage, 2012. 62(2): p. 782–90.

25. Jenkinson, M., et al., Improved optimization for the robust and accurate linear registration and motion correction of brain images. Neuroimage, 2002. 17(2): p. 825–41.

26. Woolrich, M.W., et al., Temporal autocorrelation in univariate linear modeling of FMRI data. Neuroimage, 2001. 14(6): p. 1370–86.

27. Woolrich, M.W., et al., Multilevel linear modelling for FMRI group analysis using Bayesian inference. Neuroimage, 2004. 21(4): p. 1732–47.

28. Stephan, K.E., et al., Dynamic causal models of neural system dynamics:current state and future extensions. J Biosci, 2007. 32(1): p. 129–44.

29. Havlicek, M., et al., Dynamic Granger causality based on Kalman filter for evaluation of functional network connectivity in fMRI data. NeuroImage, 2010. 53(1): p. 65–77.

30. Poch, C., et al., Time-varying effective connectivity during visual object naming as a function of semantic demands. J Neurosci, 2015. 35(23): p. 8768–76.

31. Friston, K.J., C.D. Frith, and R.S.J. Frackowiak, Time-dependent changes in effective connectivity measured with PET. Human Brain Mapping, 1993. 1(1): p. 69–79.

32. Ochsner, K.N. and J.J. Gross, The cognitive control of emotion. Trends Cogn Sci, 2005. 9(5): p. 242–9.

33. Noel, X., M. Van Der Linden, and A. Bechara, The Neurocognitive Mechanisms of Decision-making, Impulse Control, and Loss of Willpower to Resist Drugs. Psychiatry (Edgmont), 2006. 3(5): p. 30–41.

34. Paret, C., et al., fMRI neurofeedback of amygdala response to aversive stimuli enhances prefrontal-limbic brain connectivity. Neuroimage, 2016. 125: p. 182–188.

35. Nicholson, A.A., et al., The neurobiology of emotion regulation in posttraumatic stress disorder: Amygdala downregulation via real-time fMRI neurofeedback. Hum Brain Mapp, 2017. 38(1): p. 541–560.

36. Shahbabaie, A., et al., Transcranial DC stimulation modifies functional connectivity of large-scale brain networks in abstinent methamphetamine users. Brain and Behavior, 2018.

37. McHugh, M.J., et al., Executive control network connectivity strength protects against relapse to cocaine use. Addict Biol, 2017. 22(6): p. 1790–1801.

38. Goldstein, R.Z. and N.D. Volkow, Dysfunction of the prefrontal cortex in addiction: neuroimaging findings and clinical implications. Nat Rev Neurosci, 2011. 12(11): p. 652–69.

39. Shahbabaie, A., et al., Transcranial DC stimulation modifies functional connectivity of large-scale brain networks in abstinent methamphetamine users. Brain Behav, 2018. 8(3): p. e00922.

40. Yin, S., et al., Amygdala Adaptation and Temporal Dynamics of the Salience Network in Conditioned Fear: A Single-Trial fMRI Study. eNeuro, 2018. 5(1).

41. Brody, A.L., et al., Neural substrates of resisting craving during cigarette cue exposure. Biol Psychiatry, 2007. 62(6): p. 642–51.

42. Sutherland, M.T., et al., Resting state functional connectivity in addiction: Lessons learned and a road ahead. Neuroimage, 2012. 62(4): p. 2281–95.

43. Gradin, V.B., et al., Salience network-midbrain dysconnectivity and blunted reward signals in schizophrenia. Psychiatry Res, 2013. 211(2): p. 104–11.

44. Li, Q., et al., Disrupted coupling of large-scale networks is associated with relapse behaviour in heroin-dependent men. J Psychiatry Neurosci, 2017. 42(6): p. 170011.

45. Soltanian-Zadeh, S., et al. Investigating the relationship between subjective drug craving and temporal dynamics of the default mode network, executive control network, and salience network in methamphetamine dependents using rsfMRI. in SPIE Medical Imaging. 2016. SPIE.

46. Moeller, S.J. and M.P. Paulus, Toward biomarkers of the addicted human brain: Using neuroimaging to predict relapse and sustained abstinence in substance use disorder. Prog Neuropsychopharmacol Biol Psychiatry, 2018. 80(Pt B): p. 143–154.

47. Jasinska, A.J., et al., Factors modulating neural reactivity to drug cues in addiction: a survey of human neuroimaging studies. Neurosci Biobehav Rev, 2014. 38: p. 1–16.

48. Janes, A.C., et al., Brain reactivity to smoking cues prior to smoking cessation predicts ability to maintain tobacco abstinence. Biol Psychiatry, 2010. 67(8): p. 722–9.

49. Berkman, E.T., E.B. Falk, and M.D. Lieberman, In the trenches of real-world self-control: neural correlates of breaking the link between craving and smoking. Psychological science, 2011. 22(4): p. 498–506.

50. Ekhtiari, H., et al., Functional Neuroimaging Study of Brain Activation due to Craving in Heroin Intravenous Users. Iranian Journal of Psychiatry and Clinical Psychology, 2008. 14(3): p. 269–280.

51. Mokri, A., et al., Relationship between Degree of Craving and different Dimensions of Addiction Severity in Heroin Intravenous Users. Iranian Journal of Psychiatry and Clinical Psychology, 2008. 14(3): p. 298–306.

52. Azita, C., et al., Comparison of craving for opioid in opioid-dependent individuals and people under methadone maintenance treatment. Journal of Kermanshah University of Medical Sciences, 2014. 17(11): p. 718–726.

